# Practical universal *k*-mer sets for minimizer schemes

**DOI:** 10.1101/652925

**Authors:** Dan DeBlasio, Fiyinfoluwa Gbosibo, Carl Kingsford, Guillaume Marçais

## Abstract

Minimizer schemes have found widespread use in genomic applications as a way to quickly predict the matching probability of large sequences. Most methods for minimizer schemes use randomized (or close to randomized) ordering of *k*-mers when finding minimizers, but recent work has shown that not all non-lexicographic orderings perform the same. One way to find *k*-mer orderings for minimizer schemes is through the use of universal *k*-mer sets, which are subsets of *k*-mers that are guaranteed to cover all windows. The smaller this set the fewer false positives (where two poorly aligned sequences being identified as possible matches) are identified. Current methods for creating universal *k*-mer sets are limited in the length of the *k*-mer that can be considered, and cannot compute sets in the range of lengths currently used in practice. We take some of the first steps in creating universal *k*-mer sets that can be used to construct minimizer orders for large values of *k* that are practical. We do this using iterative extension of the *k*-mers in a set, and guided contraction of the set itself. We also show that this process will be guaranteed to never increase the number of distinct minimizers chosen in a sequence, and thus can only decrease the number of false positives over using the current sets on small *k*-mers.

## 1 Introduction

The minimizers technique was first used in bioinformatics to speedup the computation of read overlaps in the overlap-layout-consensus paradigm [17, 18]. In that setting, minimizers allowed reads that have a significant amount of common sequence, and therefore are likely to have an overlap, to be binned together. This quick pre-processing step avoids comparing reads that cannot have an overlap, therefore saving large amounts of computation while still guaranteeing that all matching windows will be identified.

The minimizers technique proved to be very versatile, and since this original use it has been adapted to many different applications, from *k*-mer counting [4, 10, 5], to the representation of de Bruijn graphs [1], for example for genome assembly [22], to read alignment [9, 7], metagenomics [14, 8, 21] and the sparsification of data structures [6]. This method is also used outside of the realm of bioinformatics, under the name of “winnowing”, independently introduced by Schleimer et al. [19] for detection of plagiarism. See for example Marçais et al. [12] for an overview.

The minimizers method samples a sequence by selecting the smallest *k*-mer in every window of *w* consecutive *k*-mers in the sequence (see Section 2.1 for a detailed description). By selecting *k*-mers this way, two properties are satisfied:

1. two sequences with a long exact match (i.e., a match of at least *w* + *k* − 1 bases) must select the same *k*-mers, and
2. there are no large gap between selected *k*-mers (never distant by more than *w* bases).

Applications rely on these two properties (and these properties alone) to guarantee that using minimizers gives correct algorithms. For example, the overlap computation application above relies on the first property to guarantee that two reads sharing a significant amount of sequence must be binned together, therefore avoiding false negatives (missed overlaps).

It is generally beneficial to select as few *k*-mers as possible from the sequence. For example, in the case of overlap computation, this leads to smaller bins and less computation, and in the case of sparse data structures, fewer selected *k*-mers imply a sparser data structure. The *density* is the measure of the number of selected *k*-mers over the length of the sequence (see Section 2.1) and a lower density is desirable.

The minimizers method is rather a family of methods parameterized by the length *k* of the *k*-mers, the length *w* of the windows, and the order imposed on the *k*-mer to select the smallest *k*-mer in each window. Generally the parameters *k* and *w* are constrained by the application itself. By contrast, the order on the *k*-mer is a “free” parameter: regardless of the choice of the order, the two properties above are satisfied, and the algorithm is correct for any order.

Although any choice of order leads to correct result, the order has a significant influence on the expected density of selected *k*-mers. Therefore, the choice of order with lower density leads to better performance for applications using minimizers. Finding orders with low density will improve future applications and, because any order satisfy properties (1) and (2) above, these improved orders could be retrofitted into existing applications.

The problem of finding an optimal order, i.e., an order with the lowest possible density, is still open [13]. Orenstein et al. [16] proposed a heuristic, DOCKS, that is used to create orders with low density. Unfortunately, this method has a compute time that is over exponential in *k* and is impractical for *k* ≥ 10. Even further optimizations of this heuristic which reduce computational resource requirements by approximating steps in the procedure can only be used when *k* ≤ 13. Furthermore, the use of this method requires storing a very large set of *k*-mers, too large to be practical.

Marçais et al. [13] showed a method to create optimal orders when *k* is asymptotically large. Even though these orders are close to optimal, they achieve low expected density only for values of *k* that are too large in practice.

We describe here a new method to generate orders with low expected density for practical values of *k* and *w*. The proposed method is a heuristic that generates orders for values of *k* up to and greater than 500. It uses DOCKS as a starting point and extends the order to larger values of *k* with even lower density than that obtained with DOCKS. Moreover, the representation of the order is extremely compact (storing general sets in text files sized from a few kB to a few hundred MB).

We evaluate the method in two different settings. In the first setting, the orders generated are generic and the density is low in expectation. These orders would be used in applications where the sequences are not known ahead of time. In the second settings, the sequence is known ahead time (say the human genome reference sequence). Then, our method generates orders with lower density on this particular sequence than generic orders achieve.

Orders for a wide selection of parameters *w* and *k*, generic and for the human sequence (GRCh38 [20]), are available on the github page, https://github.com/Kingsford-Group/remuval. We also provide a small and easy to use C++ library (requiring only the use of an additional header file) to make use of these orders.

## 2 Background and method overview

Our method to create orders of *k*-mers with low expected density relies on the creation of *universal sets* of *k*-mers with high *sparsity* [11]. In this section, we give proper definition of these concepts before giving an overview of the steps of the method. Section 3 gives a detailed description of the method and the proofs of the important theorems. Finally, Section 4 evaluates the performance of the method to generate orders with low density.

In the following, we consider strings defined on an alphabet Σ of size *σ* = |Σ|. A *k*-mer is a string or substring of length *k*. Given a sequence *S* ∈ Σ*, *S*[*i* : *ℓ*] denotes the substring of *S* starting at position *i* and of length *ℓ*. A window of *w k*-mers of *S* is a substring *S*[*i* : *w* + *k* − 1] that contains exactly *w* consecutive overlapping *k*-mers, namely the substrings *S*[*i* : *k*], *S*[*i* + 1 : *k*], …, *S*[*i* + *w* − 1 : *k*]. We will make the slight abuse of notation of not distinguishing between a window *ω* = *S*[*i* : *w* + *k* − 1] as a string and as a set of *k*-mers *ω* = {*S*[*j* : *k*] | *i* ≤ *j < i* + *w*}.

An order on the *k*-mers is defined by giving a rank to each *k*-mer. Formally, an order is a one-to-one function from the *k*-mers into the integers *o* : Σ^*k*^ → [1, σ^*k*^]. A *k*-mer *m*_1_ is less than *k*-mer *m*_2_ according to order *o* if its rank is less than the rank of *m*_2_: *m*_1_<_*o*_ *m*_2_ ⇔ *o*(*m*_1_) < *o*(*m*_2_).

### 2.1 Minimizers and density

We define formally the minimizers method and the main quality measure of an order, the density.

For a sequence *S*, and parameters *k*, *w* and order *o* defined on the *k*-mers, the minimizers method considers all the windows of *w* overlapping *k*-mers of *S*, that is the substrings *ω*_*i*_ = *S*[*i* : *w* + *k* − 1] for 1 ≤ *i* ≤ |*S*| − *w* − *k*. For each substring *ω*_*i*_ it finds the position of the smallest *k*-mer and adds it to the set of selected positions (use left most in case of ties). Formally, the *set of selected positions* is

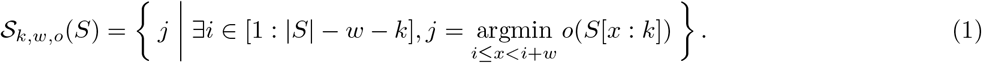

Figure 1 shows an example segment of a sequence.

**Figure 1:**
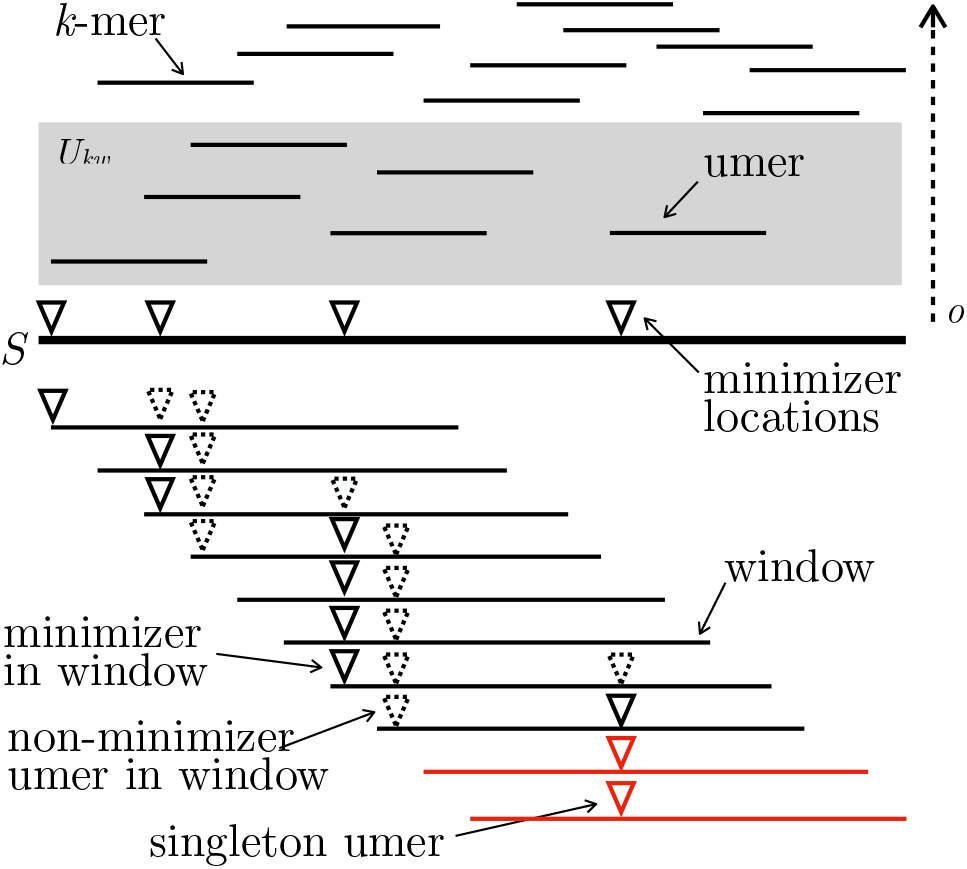
Example section of an input sequence *S* (thick black line). Above *S*, the *k*-mers of *S* drawn according to their rank for order *o*. Below *S*, the windows of *S* are represented, the triangle giving the position of the umers in the windows. One umer in this region is a singleton, meaning there is a window for which no other umer is present (in red).

As two consecutive windows *ω*_*i*_ and *ω*_*i*+1_ have almost the same *k*-mer content (they differ only in the first *k*-mer of *ω*_*i*_ and the last *k*-mer of *ω*_*i*+1_), the selected position in these two windows is likely to be identical. Therefore the minimizers method is a sampling of positions in the string *S*. The *particular density* of the sampling is the proportion of the selected *k*-mers over the total number of *k*-mers in *S*:

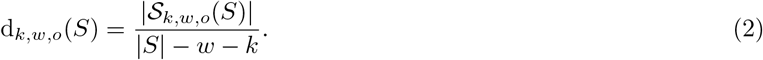

The particular density is the density of an order *o* on a given sequence *S*. The *density* d_*k,w,o*_ of order *o* (for parameter *k* and *w*) is the expected density computed, at the limit, on an infinitely long random sequence with characters chosen IID. Hence the density, defined as an expected value, is not dependent on the choice of a particular sequence.

Although the density is defined as an expected value, it can be computed exactly by computing the particular density of a specific finite sequence. A *de Bruijn sequence B*_*k*_ of order *k* is any sequence of length *σ*^*k*^ + *k* − 1 containing every possible *k*-mer exactly once [2]. As shown by Marçais et al. [11] d_*k,w,o*_ = d_*k,w,o*_(*B*_*m*_) for any *m* ≥ *w* + *k*. In other words, a de Bruijn sequence of high enough order behaves with respect to the minimizers method like an infinite sequence, and this relation allows one to compute the density for an order exactly rather than approximately.

### 2.2 Universal sets and compatible orders

*Universal k-mer sets* are central to the construction of orders with low density [11]. In fact, the proposed method, just like DOCKS [16, 15], is a heuristics to construct universal sets.

A universal *k*-mer set *U*_*k,w*_ ⊆ Σ^*k*^ is a subset of the *σ*^*k*^ possible *k*-mers that is unavoidable for any sequence of *w* consecutive *k*-mers. That is, any string of length *L* = *w* + *k* − 1, which contains exactly *w k*-mers, contains at least one *k*-mer from the set. We use simply *U* when *k* and *w* are arbitrary or can be inferred from context. The terms universal *k*-mer and *umer* (the contraction of “universal mer”) are used interchangeably to refer to a *k*-mer *u* ∈ *U*.

The set of all *k*-mers Σ^*k*^ is necessarily a universal set, but much smaller sets exists. The DOCKS heuristic is one possible heuristics to generate such sets, and we propose here another method.

Given a universal set *U*_*k,w*_, an order *o* on the *k*-mers is *compatible* with *U*_*k,w*_ if the umers always compare less than the *k*-mers not in *U*_*k,w*_. That is, for any *u* ∈ *U*_*k,w*_ and *v* ∉ *U*_*k,w*_, then *u* <_*o*_ *v*. Although there are many orders compatible with the universal set *U*_*k,w*_, we consider here any compatible order.

When using an order *o* that is compatible with the set *U*_*k,w*_ in the minimizers method on *S*, then only *k*-mers from *U*_*k,w*_ are ever selected. This holds because *U*_*k,w*_ is universal, therefore every possible window contains at least one element of *U*_*k,w*_, these elements of *U*_*k,w*_ compare less than any *k*-mer not in *U*_*k,w*_, and the smallest element of *U*_*k,w*_ in the window is selected. Hence, only the umers are relevant and the properties of the universal set define the density of a compatible order.

One characteristics of universal sets is of particular interest: the *sparsity*, denoted Sp(*U*). Every window *ω* ∈ Σ^*k*^ must contain at least one umer. Some windows may contain exactly one umer. The proportion of windows containing exactly one umer over the total number of windows *σ*^*w*+*k*−1^ is called the sparsity of the universal set. Marçais et al. [11] showed that the expected density of any order compatible with *U*_*k,w*_ is correlated with the sparsity of the universal set: the higher the sparsity of a universal set, the lower the density of a compatible order.

### 2.3 Method overview

Our method is an iterative process that starts from universal set 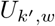 for a small length of mers (say *k*′ < *k*) and extends, one base at a time, the mers of 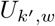 to obtain a universal set *U*_*k,w*_ for the desired mer length *k*. At each iteration, the universal set is optimized to be smaller and to increase the sparsity (and therefore a low expected density). Any universal set 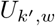 can be used as a starting point. In practice, we use the DOCKS-generated universal sets as a starting point as these sets already give compatible orders with low expected density.

There are two operations at each iteration: naïve extension and optimization (known as reM_u_val). The naïve extension (Section 3.1) is a simple procedure to create (*k* + 1)-mers from *k*-mers which has three interesting properties: first the set of extended mers is also a universal set for (*k* + 1) and *w* (Theorem 1), second the sparsity of the extended set *U* · Σ is equal to the sparsity of *U* (Theorem 2), and third the density of the compatible orders for the extended set is less or equal to the density of the original set (Theorem 3).

The second operation of reM_u_val (Section 3.2) uses an Integer Linear Program (ILP) to further reduce the size of the extended set and increases the number of *singleton umers* (see Section 3.3 for proper definition). The operation of increasing singleton umers is not directly equivalent to increasing sparsity (or lowering density), but it is related and the sparsity of the universal set is guaranteed to be non-decreasing. Additionally, the reM_u_val operation also preserves the universal property of the *k*-mer set (Theorem 4) and can only lower the sparsity (Theorem 5). The difficulty here is to have an ILP of reasonable size—that does not grow exponentially in size—and that is consequently solvable in a reasonable time in practice. Our method uses a particular *trie* to encode the umers in an efficient way, and only needs to consider a small subset of all possible windows of *w k*-mers to solve the ILP to optimality (Section 3.3).

## 3 Methods

### 3.1 naïve extension

The naïve extension of a *k*-mer set *U* is the operation of appending every letter of the alphabet to every *k*-mer from a set:

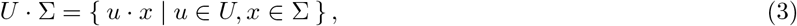

where · is the concatenation operator. The first important property of naïve extension is that it preserves universality.

#### Theorem 1.

*The naïve extension U*_*j,w*_ · Σ *of a universal set U*_*j,w*_ *is universal.*

*Proof.* Let *p* = (*m*_1_, *m*_2_, …, *m*_*w*_) be a path of *w* nodes in the de Bruijn graph *G*_*j*+1_. The de Bruijn graph of order *j* + 1 is the line graph of the de Bruijn graph of lower order *j*. Therefore, there is a corresponding path *p*′ of *w* + 1 nodes in *G*_*j*_, where each node in *p* is the edge between two nodes in *p*′. Because *U*_*j,w*_ is universal, at least one of the *w* nodes of 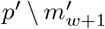, say 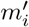, is in *U*_*j,w*_. By construction, the (*j* + 1)-mer represented by the edge 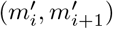 is in *U*_*j*+1,*w*_, and in turn the corresponding node *m*_*i*_ ∈ *p* is also in the set *U*_*j*+1,*w*_. Since this holds for any paths *p*′ in *G*_*j*+1_, *U*_*j*+1,*w*_ is also universal.

Naïve extension also does not change the sparsity.

#### Theorem 2.

*The sparsity of the set U* · Σ *is equal to that of U.*

*Proof.* Let *S* be a de Bruijn sequence of order *L* + 1, for *L* = *w* + *k* + 1 and let *ω* = *S*[*i* : *w* + *k* − 1] be a window of *S* that contains only one *k*-mer from *u* ∈ *U*. If *u* is at position *x* (i.e., *u* = *S*[*x* : *k*]), then the (*k* + 1)-mer *u*′ = *S*[*x* : *k* + 1] is in *U* · Σ, and no other (*k* + 1)-mer of *ω*′ = *S*[*i* : *w* + *k*] is in *U* · Σ. Hence, the sparsity of *U* · Σ is greater or equal that of *U*.

Conversely, a similar argument shows that the sparsity of *U* · Σ is less or equal that of *U*, and both sets have the same sparsity.

Naïve extension, in some sense, also does not increase the density. Because the notion of density is defined for orders and not universal sets, and because many orderings are compatible with a universal set *U*, this statement of not increasing density must be properly qualified.

Let *o*_*k*_ and *o*_*k*+1_ be orders defined on the *k*-mers and (*k* + 1)-mers respectively. Then order *o*_*k*+1_ is an *extension* of order *o*_*k*+1_ means that if *k*-mer *m*_1_ compares less than *k*-mer *m*_2_ according to *o*_*k*_, then any extension of *m*_1_ compares less than any extension of *m*_2_ according to *o*_*k*+1_. Formally, for all pairs of distinct *k*-mers *m*_1_, *m*_2_ ∈ Σ^*k*^ and for all *x, y* ∈ Σ:

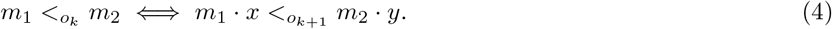

This property holds, for example, for the lexicographic orders.

#### Theorem 3.

*Given orders o and o*′ *compatible, respectively, with the universal sets U*_*k,w*_ *and its extension U*_*k,w*_ · Σ, *where o*′ *is an extension of o, then*

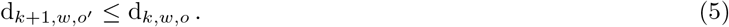

*Proof.* Let *S* be the de Bruijn sequence of order *L* + 1, for *L* = *w* + *k* − 1, i.e., a de Bruijn sequence of order large enough to compute the density for both order *o* and *o*′. Consider 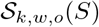 and 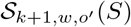, the locations in *S* that are chosen by the minimizer method for order *o* and *o*′ respectively. Because *o*′ is an extension of *o*, the set 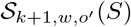 is a subset of 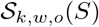.

Suppose it is not, and there exists a position 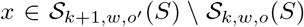. Then the (*k* + 1)-mer *m*′ at position *x* compares less than all other (*k* + 1)-mer in some window *ω*′ = *S*[*i* : *w* + *k*] (where *i* ≤ *x* < *i* + *w* + *k*). Then *m*, the *k*-long prefix of *m*′, must compare less than all other *k*-mer in the window *ω* = *S*[*i* : *w* + *k* − 1], and *x* is also in 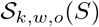. Contradiction.

It is possible for the density to decrease, and not just stay constant, by naïve extension. In the following example illustrated in Figure 2, we consider the effect of naïve extension on tie breaking in the minimizers method. Assume there is a pair of *k*-mers *x* < *y* ∈ *U*_*k,w*_ in a sequence *S* such that one instance of *x* appears to the left two instances of *y*. For example, *x* = *S*[*i* : *k*], *y* = *S*[*j*_1_ : *k*] and *y* = *S*[*j*_2_ : *k*], where *i* < *j*_1_ < *j*_2_, and that every other umer that are close to the instances of *x* and *y* (say within *w* bases) compare higher than *y* (hence they can be ignored). Then all 3 position *i*, *j*_1_ and *j*_2_ are part of the selected positions.

Suppose that after extensions, the (*k* + 1)-mers *x* = *S*[*i* : *k* + 1], *y*_1_ = *S*[*j*_1_ : *k* + 1] and *y*_2_ = *S*[*j*_2_ : *k* + 1] satisfy *x* < *y*_2_ < *y*_1_. This is possible because the two extensions of *y* may not be equal. This order is possible even for an extension of the order of *U*. Then there may not be any window where *y*_1_ is the smallest umer and only the positions *i* and *j*_2_ are selected, thereby lowering the density.

While naïve extension preserves universality and sparsity, and does not increase density, it does not preserve the optimality of any property. That is, even if *U* is a universal set of minimum sparsity or minimum density or minimum size, the naïve extension *U* · Σ is not guaranteed to be of minimum sparsity or minimum density or minimum size. This justifies the second step of optimizations to obtain a better set of *k*-mers.

**Figure 2:**
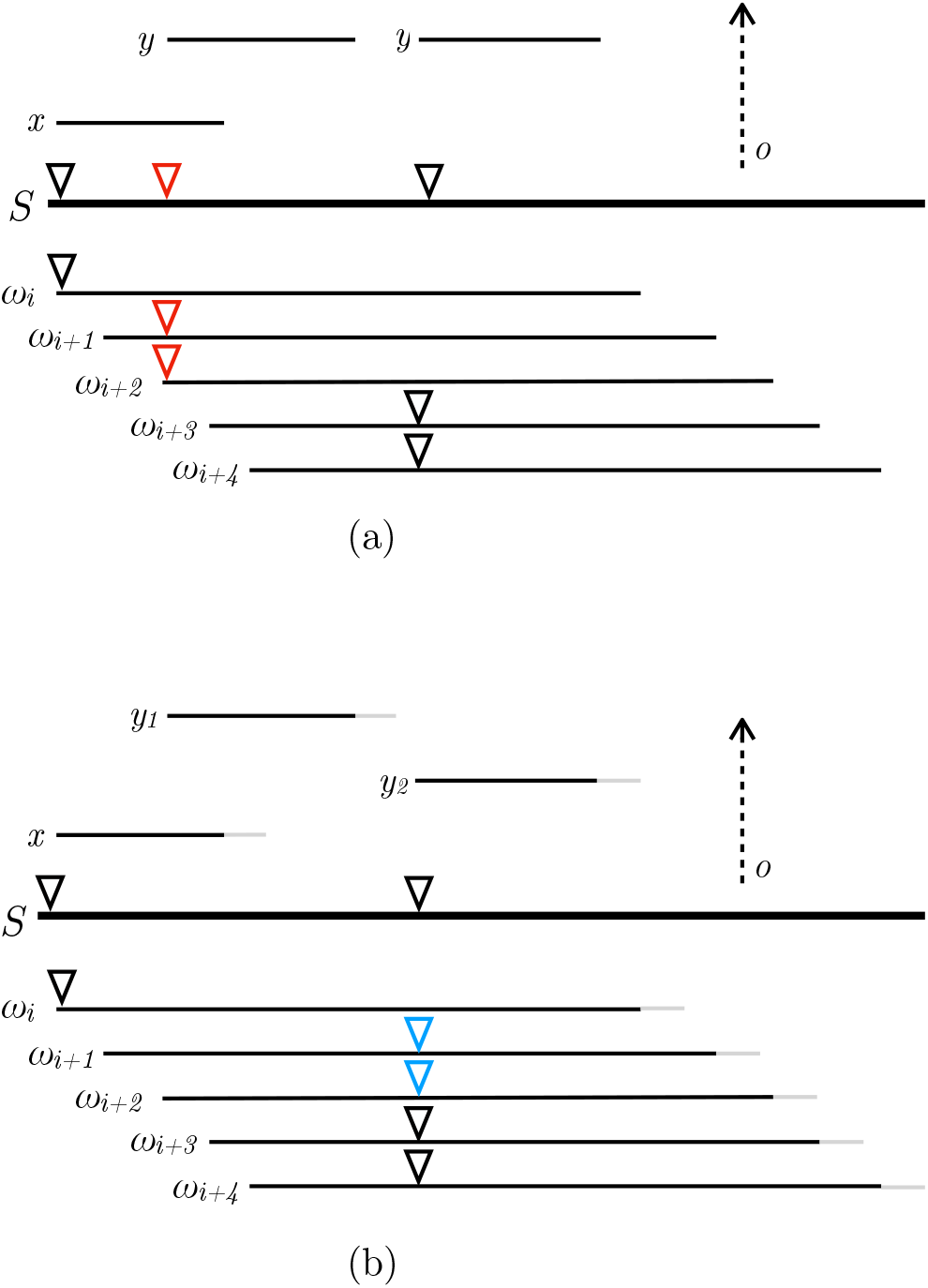
Minimizer locations on sequence *S* (a) before and (b) after naïve extension of *k*-mers *x* and *y* and windows *ω*_*i*_, …, *ω*_*i*+4_. Before extension the left most copy of *y* is the minimizer in both windows *ω*_*i*+1_ and *ω*_*i*+2_ (red triangles). After extension the two instances of *y* are different, *y*_1_ is not the minimum in any window and *y*_2_ is now the minimum in *ω*_*i*+1_ and *ω*_*i*+2_ (blue triangles). Therefore extension reduced the number of selected positions and lowered the density.

### 3.2 *M*_*u*_ and reM_u_val

The naïve extension operation generated sets that are not optimal for density. The reM_u_val operation removes elements from a universal set *U* in order to lower the density of the compatible orders.

Given a universal set *U* which is not of minimum size, there exists a *k*-mer *u* such that its removal from *U* leaves a smaller but still universal set, that is the set *U* \ *u* is also a universal set. Observe that if there exists any window *ω* of *w k*-mers for which *u* is the only element from *U* in *ω* (i.e., *ω* ∩ *U* = {*u*}), then *u* cannot be removed from the set, or *ω* would no longer contain a *k*-mer in the reduced set. Conversely, if every window that contains *u* also contains another element from *U*, we can safely remove *u* from *U* and still maintain the universality. We can formally define the minimum co-occurrence measure for an element *u* ∈ *U* as

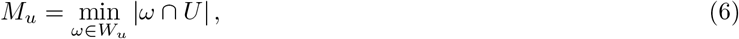

where *W*_*u*_ is the set of all windows that contain *u*.

#### Theorem 4.

*Given a universal k-mer set U and a umer u* ∈ *U such that M*_*u*_ > 1, *the set U* \ *u is also a universal set.*

*Proof.* For any window *w* that contains *u* there must exist some *k*-mer *u*′ ∈ *U* that is also in *w*, otherwise *M*_*u*_ would be 1. Therefore all windows still contain at least one *k*-mer from *U* \ *u*.

We can then remove any *u* ∈ *U* where *M*_*u*_> 1, to produce a smaller universal set. A umer with a co-occurrence value of 1 (i.e., *M*_*u*_ = 1) is called a *singleton* umer. We refer to the *removal* of the non-single umers, i.e., the umers with high co-occurrence value *M*_*u*_ > 1, from the set *U* as reM_u_val.

This operation does not decrease the sparsity.

#### Theorem 5.

*If U and U* \ *u are universal sets, then*

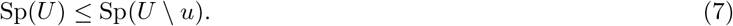

*Proof.* Any window has necessarily fewer elements from *U* \ *u* than from *U*. Hence, *U* \ *u* has at least as many windows with only one *k*-mer and its sparsity is at least as large as that of *U*.

Unfortunately, the question of whether the density of compatible orders with *U* and *U* \ *u* increases or decreases is not as simple. For example, take an order *o* such that *u* is the largest element in *U*. Then *u* is never selected as a minimizer in any window, because every window contains another element of *U* that compare less than *u*. Then the order *o* is also compatible with *U* \ *u* and the density is unchanged. On the other hand, when *u* is not the largest element according to *o*, the density could go up or down.

Even though we cannot prove a general result on how reM_u_val affects density, in practice we always see a reduction (see Section 4).

The order in which the *k*-mers are removed is significant: if *u* and *u*′ have high co-occurrence value, after removing *u*′ from the universal set, it is possible for *u* to now be a singleton in *U* \ *u*′, for example if there exists a window *ω* such that *ω* ∩ *U* = *u, u*′. It may not be possible to remove all the umers with high co-occurrence value and depending on the order of removal, the total number of removed umers may vary.

### 3.3 ILP formulation of reM_u_val

Our reM_u_val method uses an ILP to minimize the number of umers to keep while still having a universal set. We will define only one set of binary variables, *y*_*u*_, which are 1 when umer *u* is retained and 0 when it should be removed for all *u* ∈ *U*. In the following ILP, *W* = Σ^*w*+*k*−1^ the set of all possible windows:

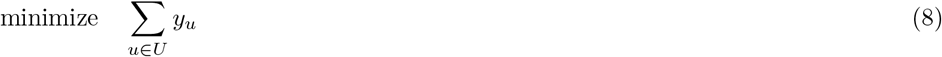

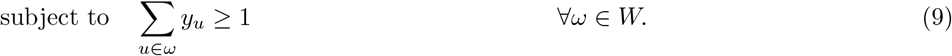

This ILP is deceivingly simple, as part of the structure of the problem is not directly encoded in the linear constraints themselves, but in the set *W*.

Although the objective function of the ILP does not explicitly attempt to maximize the number of singleton *k*-mers, after running the ILP, every *k*-mer that is kept must be a singleton. By contradiction, if there existed a umer *u* in the set *U*′ of *k*-mers kept by the ILP with a high co-occurrence value, then the set *U*′ \ *u* would still be a universal set and setting *y*_*u*_ to 0 would give a lower objective.

Also, as expressed above, the size of the ILP is exponential in *k* and *w*. Rather than recreating this ILP from scratch at each iteration of the naïve extension and reM_u_val loop, we modify the ILP from the previous iteration and use the singleton umers to reduce the size of the problem.

First, any singleton umer must be kept, hence the ILP will set *y*_*u*_ = 1. Given that these variables are not necessary they are not included in the ILP. Moreover, the set of singleton umers keeps on growing and can be partially inferred from one iteration to the next. If *u* ∈ *U* is a singleton *k*-mer, then there exists a window *ω* where *ω* ∩ *U* = *u*, say *u* = *ω*[*i* : *k*]. In every extension of the window *ω* · *x*, where *x* ∈ Σ, every (*k* + 1)-mer is an extension of a *k*-mer of *ω*. Therefore, the only (*k* + 1)-mer of *ω* · *x* that is in *U* · Σ is the extension *u*′ = *S*[*i* : *k* + 1] of *u*, and *u*′ must have a co-occurrence number of 1. Hence, at least one extension (and sometimes all extensions) of a singleton umer is also a singleton umer.

In practice, these singleton umers are kept in a modified trie [3]. This implementation is efficient as when all the extensions of a singleton are also singletons, it takes no extra memory to store the extensions.

Second, any window which contains a singleton umer necessarily satisfies the constraint in eq 9, regardless of the values of the other variables. Therefore, this constraint needs not be included in the ILP. Also, the windows that will be included in the ILP at the next iteration is a subset of the extensions of the windows that are considered in the current iteration. The algorithm keeps track of only these windows.

The two remarks above allows to construct the ILP without having to consider all possible windows, and also notably reduces the size of the problem to solve.

### 3.4 Sequence-specific reM_u_val

Universal *k*-mer sets are guaranteed to always have one *k*-mer in each *w*-long window, but many application that use minimizer schemes will only be searching over a single reference sequence, for instance the human genome. For example, in a read to genome aligner, where the index of the genome is fixed, it is legitimate to optimize the minimizers scheme for the particular genome sequence.

As *k* increases, the fixed reference sequence does not contain all possible windows and thus some of the universal mers cannot occur as minimizers. In these cases the set of *k*-mers that is used to define a minimizer scheme can be reduced even further to create a new set 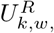, which is only universal for the reference sequence *R*. This new *k*-mer set is not universal for all strings, but if a window is found that does not contain a *k*-mer from *U*^*R*^ we are guaranteed that this window is not contained in *R*. This guarantees, for example, that that even though the set *U*^*R*^ is not universal, the above application of alignment to a genome is still correct: a read not containing a minimizers from *U*^*R*^ does not align to the genome *R*.

We call “sequence-based reM_u_val” (described below) the operation restricting a universal set to a specific sequence. There are two possible ways to generate sequence specific universal sets. Either by applying sequence-based reM_u_val first or applying it last. That is, either (1) run the reM_u_val and extension loop to construct *U*_*k,w*_ then run sequence-based reM_u_val to restrict that set, or (2) run sequence-based reM_u_val to obtain 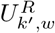 then run an extension and sequence-based reM_u_val loop. These procedures give similar results but, maybe surprisingly, the second method is less computationally intensive (see Section 4.2).

#### Sequence-based reM_u_val

Consider a reference sequence *R*. Given a universal set *U*′ (either a general *U*_*k,w*_ or some 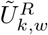 constructed by naïve extension), a reference restricted *k*-mer set 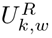 is constructed by including in the ILP (eq 9) only the windows of *R* that contain more than one umer (element of *U*′). While some umers in *U*′ are universal singletons, they may not be *sequence-specific* singletons, and these must be identified in order to reliably leave out any window from the ILP. To do so we scan the reference sequence at each iteration before the reM_u_val operation.

## 4 Evaluation of reM_u_val to find universal sets for large *k*

We use three metrics of universal *k*-mer set quality to compare various methods of construction:

- **Size** – the number of *k*-mers chosen to be in the universal set, as the fraction of the entire set of *k*-mers.
- **Density** – using a minimizer set that is consistent with the universal set, and lexicographic for all umers. (see Section 2.1)
- **Sparsity** – the fraction of windows with only one umer. (see Section 2.2)

We will compare these results with a random ordering of all *k*-mers, which a subset of the authors found to be a good proxy to many other methods in previous work [11], and when appropriate with the randomized minimizer ordering over all *k*-mers. Throughout this sections we use a constant window length *w* = 6 for comparison.

### 4.1 Universal *k*-mer sets

#### Size

Figure 3 shows the universal *k*-mer set size of our procedure as the *k*-mer size is increased. Each of the points on the plot shows the number of *k*-mers of a universal *k*-mer set as a fraction of the total number of *k*-mers. The sets were constructed by starting with DOCKS sets of various *k*′-mer lengths (*k*′ < *k*), and are compared to the sizes of the initial DOCKS sets. The size of the final set size is somewhat dependent on the size of the set when reM_u_val is started. This is likely due to the fact that “bad” choices being made early cannot be corrected. That is, once one *k*-mer is alone in a window at least some of its extensions must be in all future sets. While DOCKS has more freedom to not be constrained in the greedy framework, it comes at the expense of the computational resources, and as we can see, our method is able to construct sets for larger values of *k*.

**Figure 3:**
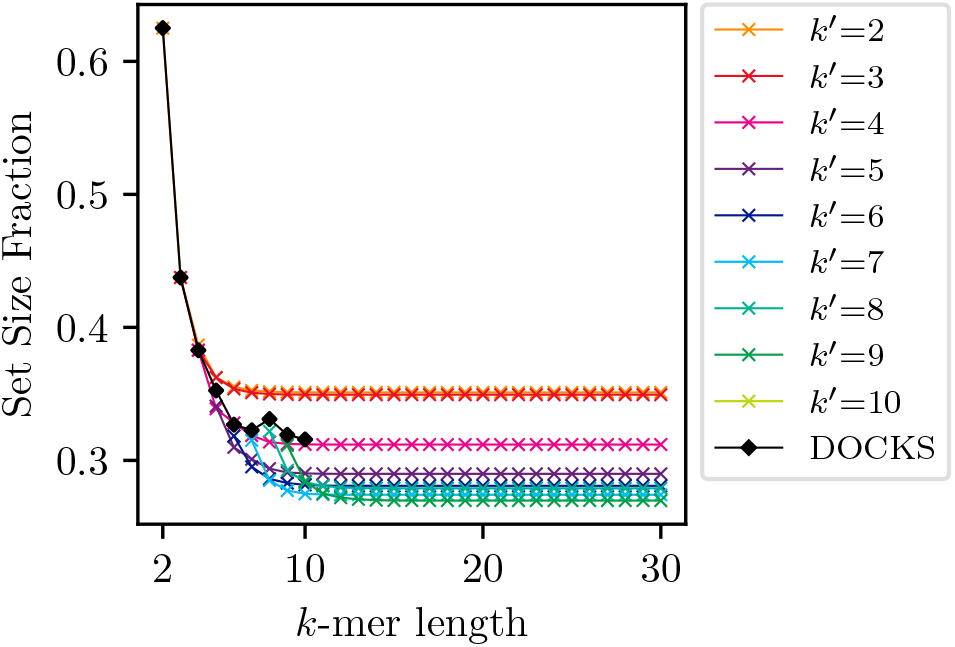
Universal *k*-mer set relative sizes for naïve extension and reM_u_val with various starting sequence lengths, *k*′, compared to that of DOCKS. The set size fraction decreases with each iteration of the iterative procedure, and is always lower than that of the DOCKS set from which the procedure started started.

#### Density

As described in Section 2.1 the expected density (and sparsity) for an ordering can be calculated using a de Bruijn sequence of a long enough order. While this is useful in understanding the performance of an ordering in general, it is not always feasible in practice. For instance, the de Brujin sequence of order 21 is already ≈ 4 TB for the DNA alphabet, and is increased in size approximately σ-fold for each additional base in the mer length. Therefore, while we are able to calculate universal sets for *k* > 500, we can only compute the expected density up to *k* ≤ 15.

Figure 4 shows the densities of a minimizers that are compatible with universal *k*-mer sets constructed using various methods across various values of *k*. There is no obvious relationship between the density of the initial DOCKS and the eventual asymptotic density of extension procedure. While the question of guaranteeing that density will not increase after reM_u_val is still open, it is clear that in practice this never happens. Quite often the density after the initial rounds of reM_u_val is decreased substantially. In fact, the reM_u_val procedure is able to decrease density of the DOCKS sets even before the first round of naïve extension.

**Figure 4:**
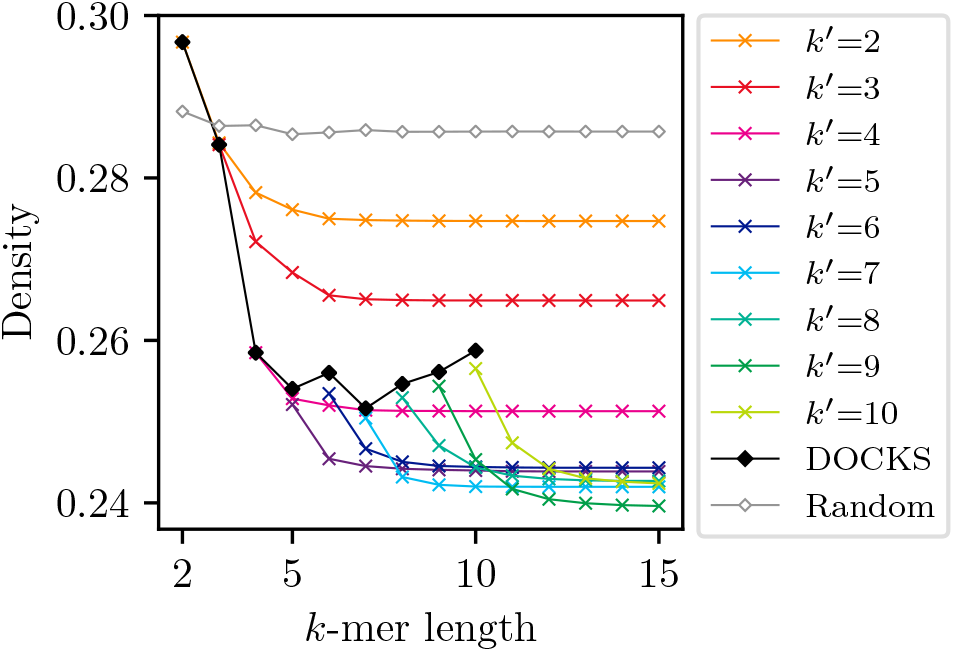
Universal *k*-mer expected density for naïve extension and reM_u_val with various starting sequence lengths, *k*′, compared to that of DOCKS. The density decreases with each iteration of the iterative procedure, and is always lower than that of the DOCKS set from which the procedure started started.

#### Sparsity

While density and sparsity are negatively correlated, the exact relationship is complicated: the sparsity is a property of the universal set and the density is a property of a particular compatible order. We additionally show in Figure 5 the sparsity of the same universal sets shown in Figure 4. These results confirm Theorem 5, that after reM_u_val the sparsity can only increase. While the reM_u_val increases the number of singleton *k-mers* rather than singleton *windows* (windows with only one umer, i.e., the sparsity), it is intuitive that these two measures would be linked: for each *k*-mer for which *M*_*u*_ is reduced to 1, at least one additional window also now has only one umer.

**Figure 5:**
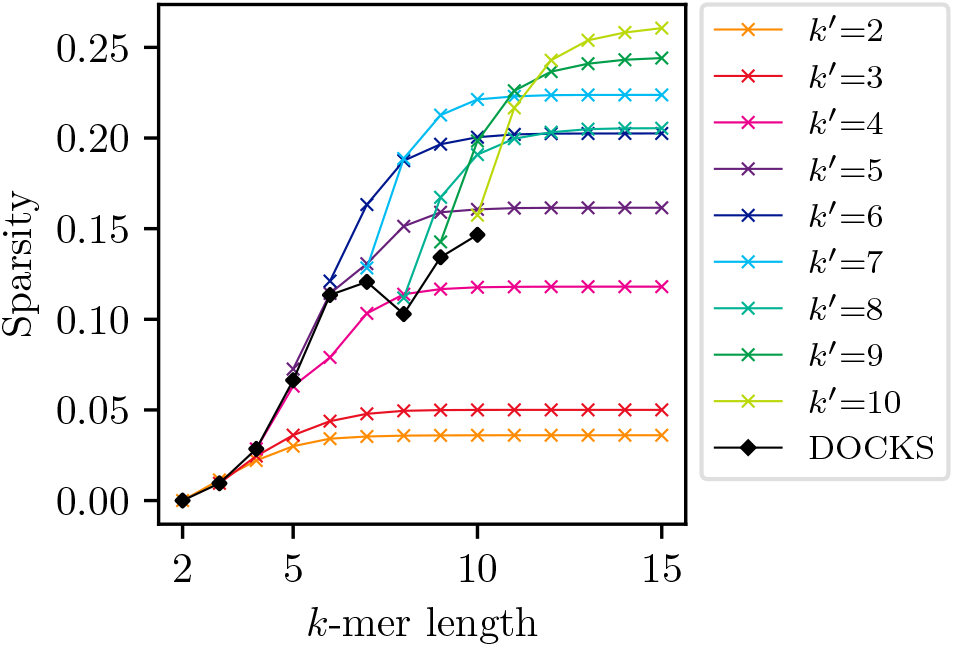
Universal *k*-mer expected sparsity for naïve extension and reM_u_val with various starting sequence lengths, *k*′, compared to that of DOCKS. The sparsity increases with each iteration of the iterative procedure, and is always higher than that of the DOCKS set from which the procedure started started.

### 4.2 Human-specific *k*-mer sets

While we present two methods for finding sequence-specific *k*-mer sets, contraction of existing sets for a given *k* and sequence-specific expansion and reM_u_val, we found that in practice the latter worked much better. In our tests, both methods achieved similar results (for *k*′ = 10, *k* ∈ [10, 17], difference in density 10^−4^, difference in sparsity < 10^−4^, difference in set size fraction < 10^−3^) but the sequence-specific iteration was much more computationally efficient.

This is mostly due to the list of windows that need to be considered in the ILP. This list is quite large when this is constructed from a universal set for large *k*. Many fewer of the *k*-mers in the universal set have a *M*_*u*_ value of 1 when only the human sequence is considered, therefore many more windows need to be included in the ILP. Consider a subset of umers with the same prefix of length *k*′ < *k* that are to be removed when we produced 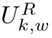 from *U*_*k,w*_. If instead we had constructed 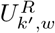 then performed expansion, the prefix of these *k*-mers would have been removed earlier.

The results in this section examine the quality of sequence-specific *k*-mer sets produced for the human reference genome (GRCh38), and compare them to universal *k*-mer sets. The sequence used additionally includes the human genome reverse complement sequence in order to include all possible windows that may be encountered when applied. As mentioned above, because iterative sequence-specific reM_u_val is more efficient than contraction of existing sets and provides similar results this is the method used throughout. Because of the computational issues that sequence-specific set construction presents only *k* ≤ 18 are presented, but the increase in quality is still considerable.

#### Size

Figure 6 shows the sizes of both universal *k*-mer sets and human-specific *k*-mer sets for increasing values of *k*. The solid lines epresent the sequence-specific sets, and the dashed lines are for the universal sets. The blue dot-dash line shows the fraction of all *k*-mers that are contained in the human genome. For values of *k* ≤ 12, the human genome sequence contains all possible *k*-mers, so the universal and human specific *k*-mer sets are approximately the same size. As the fraction of *k*-mers present in the human genome drops, so does the size of the sequence-specific *k*-mer set. We find that even though the sizes of the *k*-mer sets are much smaller for the human specific sets, the sizes of the trie needed to store them is much larger, 12 Gb vs 84 Mb for *k*′ = 10, *k* = 17. This means the data structure is less able to compress the *k*-mers in the set.

**Figure 6:**
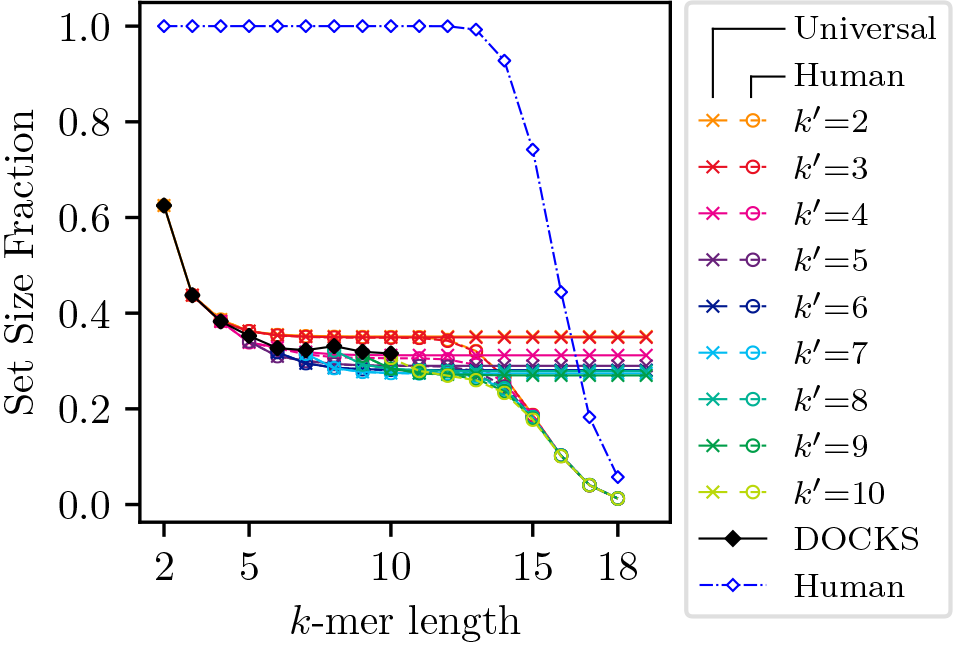
The relative size of human-specific and universal *k*-mer sets. The blue diamond indicate the fraction of *k*-mers that are present in the human genome. Human-specific are smaller than universal sets for values of *k* when the human genome does not contain all *k*-mers.

#### Density and Sparsity

Figures 7 and 8 show a decrease in the density factor and increase in sparsity of using sequence specific sets versus universal *k*-mer sets. The density factor using the human genome sequence is substantially higher than the expected density factor shown in the previous section. While it seems intuitive that the number of *k*-mers in the set would decrease as the fraction of *k*-mers in the human sequence decreases, the difference in sets in fact happens when the fraction of *windows* decreases (i.e., *k* + *w* − 1 ≥ 12) though the differences are not substantial until *k* is larger.

**Figure 7:**
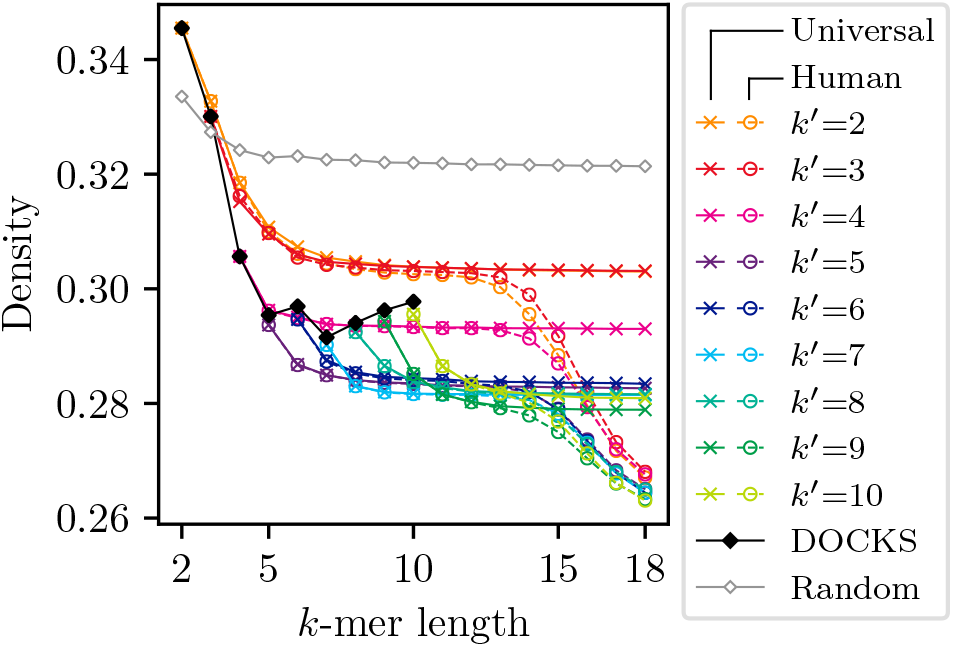
Human-specific and universal *k*-mer set densities on the human genome sequence. Generally human-specific sets have lower density than universal *k*-mer sets for larger values of *k*.

**Figure 8:**
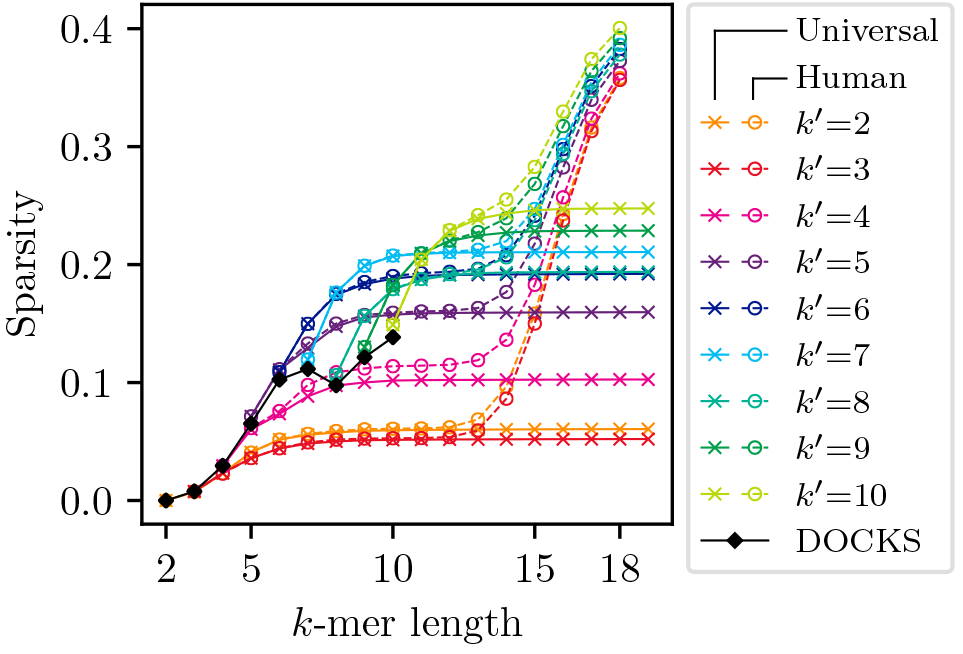
Human-specific and universal *k.*-mer set sparsities on the human genome sequence. Human-specific sets have more sparse windows than universal *k*-mer sets for larger values of *k*

## 5 Discussion and Conclusion

### DOCKS comparison

DOCKS and the method presented here are two ways to generate small universal sets with high sparsity, and these sets can then be used to obtain minimizer methods with low density. The two heuristic methods differ in their approach to generate these sets. DOCKS starts with a minimum decycling set: a set of *k*-mers that if is unavoidable by any “infinitely” long path (i.e., every cycle of the de Bruijn graph). It then iteratively expand that sets until it is unavoidable by any *w*-long path. Because it has to enumerate many paths in the full de Bruijn graph of order *k*, the methods is computationally and memory expensive for large *k*. And the lower *w* is, the more iterations are needed.

Our method starts a universal set for *k*′ and *w*, and then iteratively expand that set to *k* and *w* (*k*′ *k*). After the first iteration, only a subset of the de Bruijn graph of order *k*′ + *w* needs to be considered. When *w* is not too large, the method can create universal sets for relatively large *k*.

These two methods are complementary as they generate universal sets for different region of the parameter space. The orders compatible with these universal sets outperform, both in expectation and on the human sequence, the usual “random” order. But the gap between the best known lower bound on density and the best known orders is still wide. The development of new methods to generate orders for minimizers computation remains an interesting problem with potentially good improvement for bioinformatics algorithms.

### Larger *k* and *w*

Some applications use minimizers with large values of both *k* and *w*, for instance the default settings of MashMap[7] use a minimizers scheme with *k* = 16 and *w* = 111, which is outside the limits of both of the DOCKS and our method. Nevertheless, the universal sets generated may still be of use in that context. First, the trie data structure (see Section 3.3) used in our method can detect that a query string has a prefix in the trie. This implies that naïve extension is implemented at no computational or storage cost. Hence the sets are directly usable for larger values of *k*. Second, a set which is universal for *k* and *w* is also universal for *k* and *w*′, for any *w*′ ≥ *w*. Thus any of the sets from either DOCKS our method can be used in practice with larger *w*, albeit with a loss of performance.

### Increasing *w*

For large values of *w* our method becomes computationally intractable, both because the initial set of windows to construct the first ILP becomes very large and also because a smaller percentage of those windows can be excluded from the subsequent ILPs because they contain singleton umers. For instance, at *k* = 11 the number of windows that contain more than one umer is 138,580 for *w* = 6 but grows to 1,138,888,351 for *w* = 10 (starting length of *k*′ = 6). This means that new insights are needed in order to continue progress in generating universal sets where both *w* and *k* are large.

One possible avenue is to perform an iterative extension not only in *k*, as we presented here, but also in *w*. That is, instead of starting with a small *k*′ and *w* already at its final value, is it possible to start with small *k*′ and *w*′ and conjointly increase both values?

Keeping with the same framework of extension followed by optimization, extending *w* does not preserve the same properties as extending *k*. For example, with naïve extension, at least one of the extension of a singleton umer is also a singleton umer. This does not hold when *w* is extended: a new umer may be introduced into a window with previously only one umer. Different rules to maintain the optimization problem between iterations are necessary.

### Plateau

As discussed in Section 4, our method shows a plateau in improvement after some number of iterations. This is due to the greedy nature of the extension and the overwhelming effect of singleton umers. After at most *w* extensions of a singleton umers, all possible *σ* extensions, and not just 1, of a singleton umer are also singleton umers. Consequently, there is an exponential number of singleton umers that cannot be optimized by the reM_u_val operation, and these singleton umers create the plateau.

We can see the results of this in all of the figures above, when the initial universal set *k*-mer size is small (i.e *k*′ ≤ 3) poor choices of singleton umers can be made early limiting the quality of the subsequent solutions. One possible solution would be to extend in both directions (to the left and right of the window and *k*-mer). This would likely still provide the same plateau but possibly at a value of *k* that is twice as far.

### Sequence specific

A surprising result is that the sequence-specific *k*-mer sets are both more computationally intensive to construct and are much harder to store. This is again likely due to the singleton *k*-mers, but for the opposite to the plateau behavior mentioned above. In the generic case, the exponential set of singleton umers are very easy to encode and reduce the size of the ILP significantly but limit the performance of the method. Because a specific sequence may not contain all the possible windows after a naïve extension, there are much fewer singleton umers, which makes the ILP larger but allows for greater optimization. An open question is how to find a sequence-specific *k*-mer set that is easy to compute and/or store.

## Acknowledgments

The authors would like to thank the organizers of the Internship in Biomedical Research, Informatics, and Computer Science (iBRIC) at University of Pittsburgh, and Heewook Lee for their support of this work.

